# BigBrain 3D atlas of cortical layers: cortical and laminar thickness gradients diverge in sensory and motor cortices

**DOI:** 10.1101/580597

**Authors:** Konrad Wagstyl, Stéphanie Larocque, Guillem Cucurull, Claude Lepage, Joseph Paul Cohen, Sebastian Bludau, Nicola Palomero-Gallagher, Lindsay B. Lewis, Thomas Funck, Hannah Spitzer, Timo Dicksheid, Paul C Fletcher, Adriana Romero, Karl Zilles, Katrin Amunts, Yoshua Bengio, Alan C. Evans

## Abstract

Histological atlases of the cerebral cortex, such as those made famous by Brodmann and von Economo, are invaluable for understanding human brain microstructure and its relationship with functional organization in the brain. However, these existing atlases are limited to small numbers of manually annotated samples from a single cerebral hemisphere, measured from 2D histological sections. We present the first whole-brain quantitative 3D laminar atlas of the human cerebral cortex. This atlas was derived from a 3D histological model of the human brain at 20 micron isotropic resolution (BigBrain), using a convolutional neural network to segment, automatically, the cortical layers in both hemispheres. Our approach overcomes many of the historical challenges with measurement of histological thickness in 2D and the resultant laminar atlas provides an unprecedented level of precision and detail.

We utilized this BigBrain cortical atlas to test whether previously reported thickness gradients, as measured by MRI in sensory and motor processing cortices, were present in a histological atlas of cortical thickness, and which cortical layers were contributing to these gradients. Cortical thickness increased across sensory processing hierarchies, primarily driven by layers III, V and VI. In contrast, fronto-motor cortices showed the opposite pattern, with decreases in total and pyramidal layer thickness. These findings illustrate how this laminar atlas will provide a link between single-neuron morphology, mesoscale cortical layering, macroscopic cortical thickness and, ultimately, functional neuroanatomy.

## Introduction

The cerebral cortex has six cytoarchitectonic layers that vary depending on cortical area [1], but cannot readily be resolved using *in vivo* MRI techniques [2]. Nevertheless, cortical microstructure underpins the functional, developmental and pathological signals we can measure *in vivo* [3,4]. Thus bridging the gap between micro scale structural measurement and whole brain neuroimaging approaches remains an important challenge. To address this, we sought to create the first whole brain, 3D, quantitative atlas of cortical and laminar histological thickness.

Cortical thickness is one widely used marker of both *in vivo* and *ex vivo* cortical structure [5–7]. Early histological studies noted marked interareal thickness differences on *post mortem* histological sections [1,8], which have since been replicated [5,9] and extended using *in vivo* MRI [10], and alterations in these patterns may be seen in neuropsychiatric illness [11–13].

MRI approaches have demonstrated patterns of cortical thickness relating to functional and structural hierarchical organisation across sensory cortices of both macaques and humans [10]. While classical studies of cortical histology also observed that primary sensory regions are thinner than their surrounding secondary sensory cortices [1,8], the thickness gradients identified in MRI extend far beyond neighbouring secondary areas into association cortical areas, while such a pattern has not been systematically studied in *post mortem* brains. However MRI thickness is known to be impacted by the degree of cortical myelination [6,14], and cortical myelination exhibits similar gradients, with primary sensory areas being more heavily myelinated than secondary sensory areas [15]. Thus it remains unclear whether thickness gradients found in MRI are artefactual, driven by gradient differences in cortical myelination causing systematic cortical reconstruction errors, or truly represent the underlying histology.

Creating a cortical layer segmentation of the BigBrain, a 3D histological model of the human brain [16], offers a solution to these problems and allows us to create a link between laminar patterns and standard MRI measures. Using this dataset, we can determine whether cortical thickness gradients are evident in measurements made with much greater spatial resolution. It opens the possibility to study whether similar cortical thickness gradients are present in fronto-motor cortices such as those identified in *in vivo* neuroimaging [17]. Going beyond overall cortical thickness, it becomes possible to examine which cortical laminae contribute to these thickness gradients, enabling better characterisation of cortical structure and the potential to link these macroscale thickness gradients to changes in laminar cortical connectivity in sensory and motor hierarchies.

Sensory processing hierarchies describe the concept that the cerebral cortex is organised with gradually changing structural and functional properties from primary sensory areas, to secondary sensory areas and ultimately higher-order association areas. Multiple measurement modalities converge on similarly ordered patterns including increasing dendritic arborisation in of pyramidal neurons [18] and electrophysiological characteristics [19], laminar connectivity patterns of projecting cortical neurons [20–22], laminar differentiation [23,24], MRI cortical thickness [10], MRI myelination [15], receptor densities [25] and temporal dynamics [26]. Topographically, hierarchies are organised such that progressively higher cortical areas are located with increasing geodesic distance across the cortical surface from their primary areas [10,27]. Ordering cortical areas along these gradients provides a framework for quantifying and understanding the relationships between cortical topology, microstructure and functional specialisation.

Carrying out analyses of histological thickness gradients poses several methodological challenges. First, thickness measurements carried out in 2D are associated with measurement artefacts due to oblique slicing [28] and stereological bias. Second, manual measurement is associated with observer dependent variability, estimated to be up to 0.5 mm [8]. Third, due to the labour-intensive nature of histological analysis, many histological atlases have a small number of sample points, with studies commonly restricted to measuring around 100 cortical samples [8,29]. These factors hinder the ability to detect and map potentially subtle cross-cortical variations in cytoarchitecture as well as overall and laminar thicknesses. BigBrain offers a unique dataset to resolve histological cortical layers comprehensively in 3D, thereby providing a concrete link between microscale patterns of structure and *in vivo* markers.

We therefore set out to automate segmentation of cortical layers in 3D in order to characterise patterns of cortical and laminar thickness across sensory and fronto-motor cortical areas. To do this we used a convolutional neural network to segment profiles of histological intensity sampled between the pial and white matter. Training profiles were generated from examples of manually segmented layers on cortical regions from 2D histological sections of the BigBrain dataset. The trained network was used to segment intensity profiles derived obliquely through the 3D histological volume and generate mesh segmentations of six cortical layers. These surfaces were used to calculate cortical and laminar thicknesses. Geodesic surface distance from primary visual, auditory, somatosensory and motor areas were calculated and used as a marker of hierarchical progression. Cortical and laminar thickness gradients were calculated for each system.

## Results

The automatically identified cortical layers closely follow bands of intensity within the BigBrain (Figure 1) and continue to follow the same features beyond the limits of training examples (Figure 2a).

**Figure 1.**
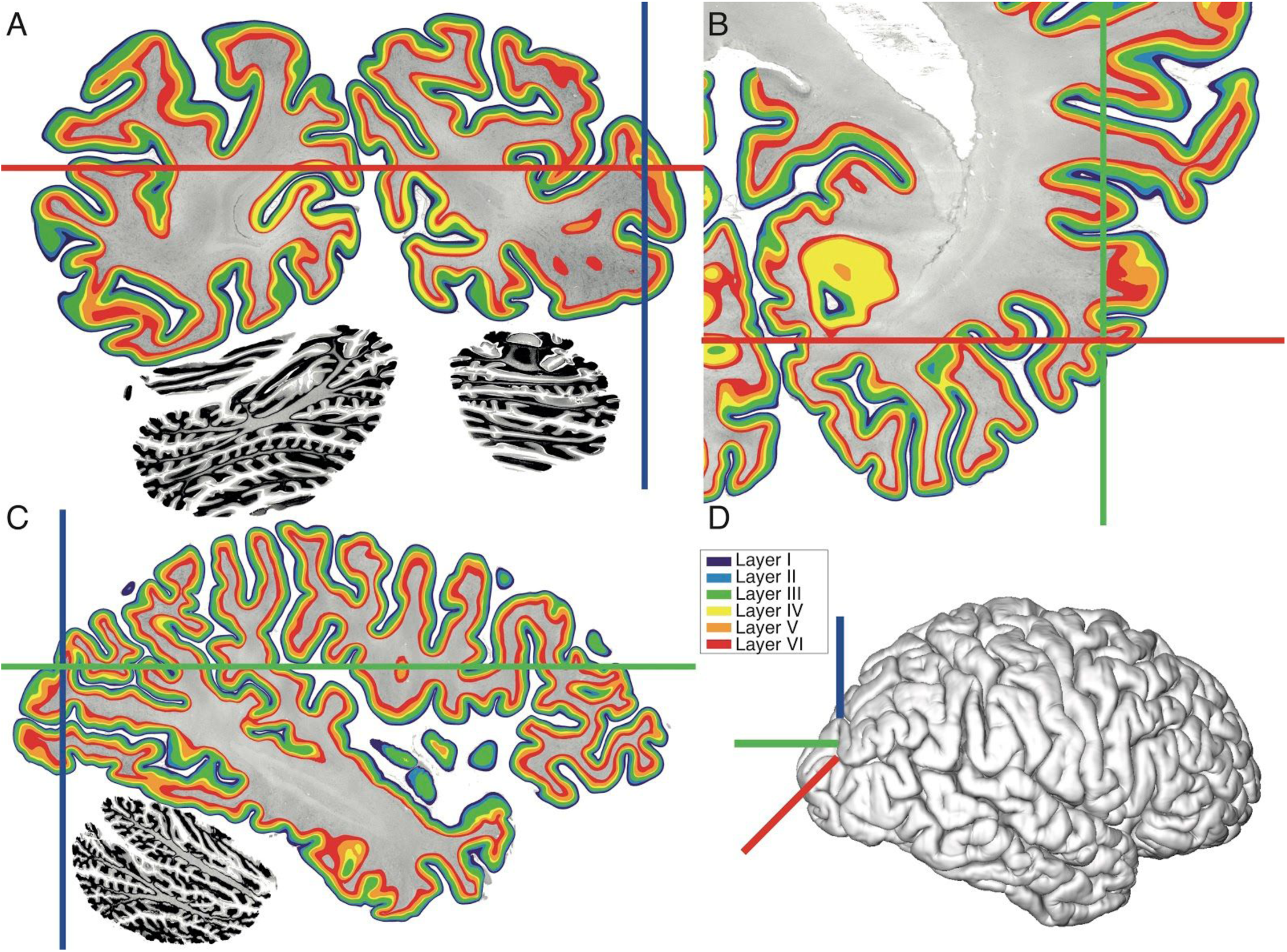
Cortical layers in 3D. Six cortical layers segmented on the 3D volume on three orthogonal planes: A=coronal, B=axial, C=sagittal. Panel D shows the location of the sections on the reconstructed pial surface of the 3D BigBrain. A, the coronal plane is the original plane of sectioning. Within this plane the axes are centered on an area of the cortex where layers would be impossible to segment in 2D, because the section only shows part of the gyrus and most layers are not visible due to the oblique sectioning of the cortex with respect to the gyrus.

**Figure 2.**
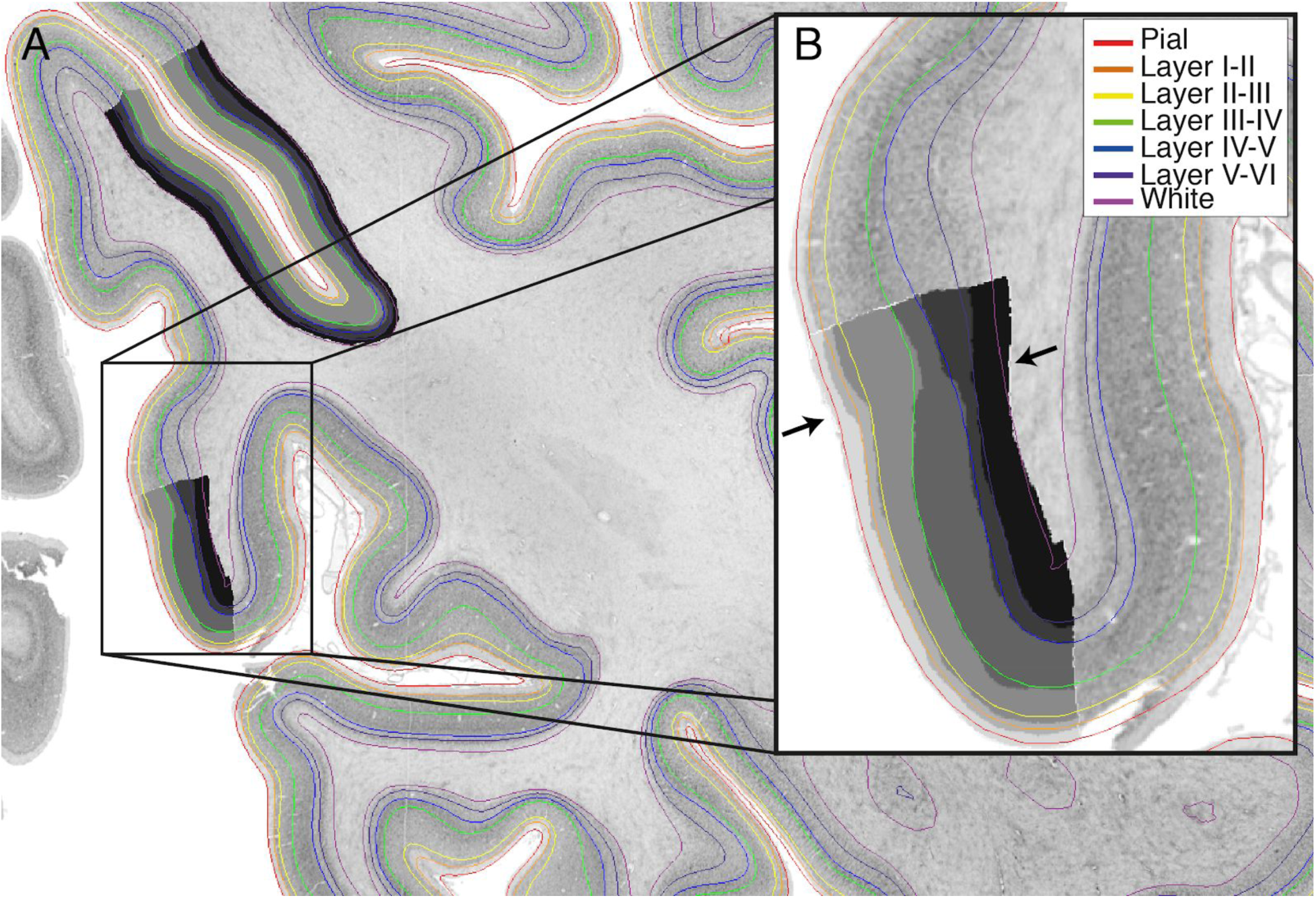
Cortical layers (coloured lines) intersected on a 2D coronal section of the right occipital cortex with manually segmented layers (superimposed grey-scale masks). A) The boundaries follow the same contours as delineated by the manually segmented training areas, and appear to accurately follow the layer bounds outside of each training area. B) At the V1-V2 boundary (marked with arrows) the thickness of layer IV changes dramatically in both manual and automated segmentations (between green and blue lines), with additional peaks in V1 intensity due to the sublayers of layer IV. As each profile is individually segmented by the network, without reference to the neighbouring profiles, the network is able to apply area-specific rules according to the shape of the profile, suggesting it might be internally identifying the area from which the profile is extracted as being either V1 or V2.

In the original BigBrain surfaces, as with MRI white matter surfaces, the white surface was placed at the maximum intensity gradient between grey matter and white matter [30]. By contrast the neural network is trained on examples where the white boundary has been manually located according to the presence of cortical neurons. This has caused a systematic shift in the location of the new white matter surface. On closer inspection, the maximum gradient at which the original surfaces were placed appears to be at the border between sublayers VIa and VIb, where the change in neuronal density is sharper than at the boundary between white matter and layer VI (Supplementary Figure 4).

A second feature apparent on visual inspection is segmentation of the layers cannot follow a single set of rules applied indiscriminately - laminar segmentations vary between cortical areas. This is most readily apparent at the V1-V2 boundary where layer IV changes considerably (Figure 2b). Layer IV is particularly broad in V1 and has multiple sublayers creating extra peaks and troughs in the intensity profiles, whereas in V2 it is much thinner and no longer differentiated into sublayers. The transition from a thick layer IV to a thin layer IV occurs precisely at the boundary between these two regions, suggesting that the network is also internally learning certain areal properties.

### Comparison of total and layer thickness maps

On visual inspection, maps of BigBrain cortical thickness are consistent with classical atlases of histological thickness reported by von Economo and Koskinas (Figure 3). In particular, the precentral gyrus is the thickest part of the cortex, with values over 4.5mm (when adjusted for shrinkage in BigBrain) and 3.5 - 4.5mm in von Economo (area FA). The thickness of the motor cortex is often underestimated in MRI thickness measurement [31], probably due to the high degree of intracortical myelination which affects the grey-white contrast, causing the white matter surface to be too close to the grey surface, such that cortical thickness is underestimated [10,14]. The calcarine sulcus is especially thin on both BigBrain (1.67 - 2.86 mm, 95% range) and von Economo (1.8 - 2.3 mm, area OC). This is also consistent with measurements from Amunts [32] of 1.47±.24mm (left) and 1.57±0.41 (right). Overall, regional values from BigBrain were highly correlated with their corresponding values in von Economo and Koskinas (left hemisphere: r=0.84, right hemisphere: r=0.83). In addition, folding-related differences are clearly visible on the BigBrain, with sulci being thinner than their neighbouring gyri.

**Figure 3.**
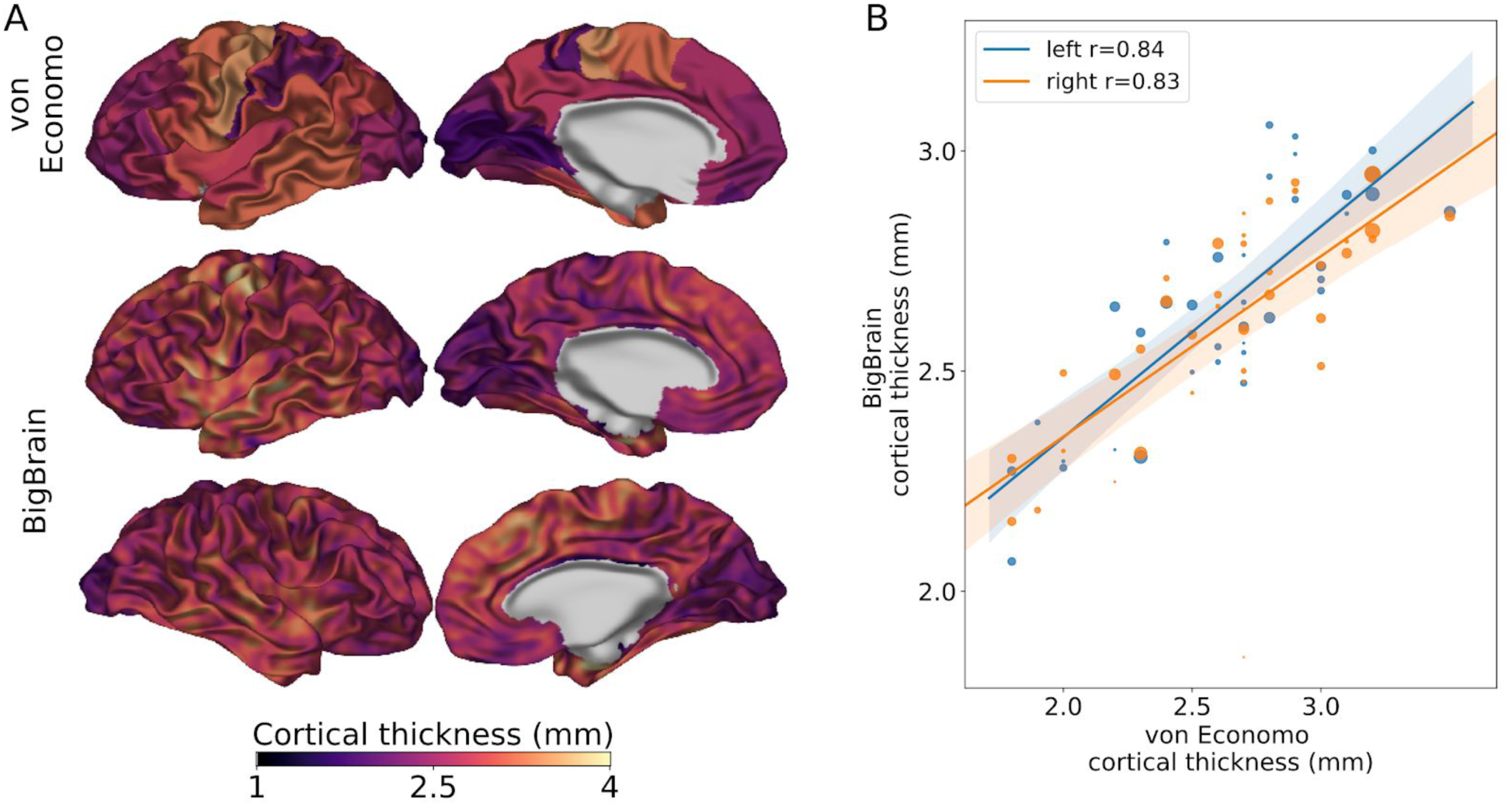
Comparison of von Economo’s maps of cortical thickness (coregistered with and visualized on the left hemisphere of BigBrain) with cortical thickness of the BigBrain for left and right hemispheres. Thickness values range from 1.8mm in the calcarine sulcus to 4.5 mm in the precentral gyrus. A) The pattern of cortical thickness across the BigBrain (displayed on smoothed surfaces, values were smoothed 3mm FWHM) matched that measured by von Economo and Koskinas. In particular, the precentral gyrus, containing the primary motor cortex, which is often underestimated with MRI, was the thickest area. Additionally, the occipital cortex around the calcarine sulcus was particularly thin in both BigBrain and von Economo. While broad trends are consistent between the two maps, BigBrain exhibits more local variability, for instance due to cortical folding, and due to the higher density of measurements made. Von Economo reported thickness measurements from around 50 cortical areas, whereas the thickness of around 1 million vertices has been measured on BigBrain. B) Regional BigBrain thickness values were highly correlated with measurements from von Economo and Koskinas. The size of each point is proportional to the area of the cortical region and overall correlations were weighted according to these areas.

BigBrain layer thickness maps are also consistent with layer thicknesses from the von Economo atlas (Figure 4). Layer III is thick in the precentral areas, and particularly thin in the primary visual cortex. Layers V and VI are thicker in frontal and cingulate cortices, but also thin in the occipital cortex. Each layer is strongly correlated with the corresponding von Economo measurements except layer II (Layer I, left r=0.48, right r=0.56, layer II, left r=-0.06, right r=-0.16, layer III, left r=0.50, right r=0.42, layer IV, left r=0.71, right r= 0.65, layer V, left r=0.63, right r=0.65, layer VI, left r=0.64, right r=0.61). The reason for the lack of agreement between layer II measurements is likely due to the low degree of intra-areal variance coupled with sources of noise such as interindividual differences and imperfect coregistration.

**Figure 4.**
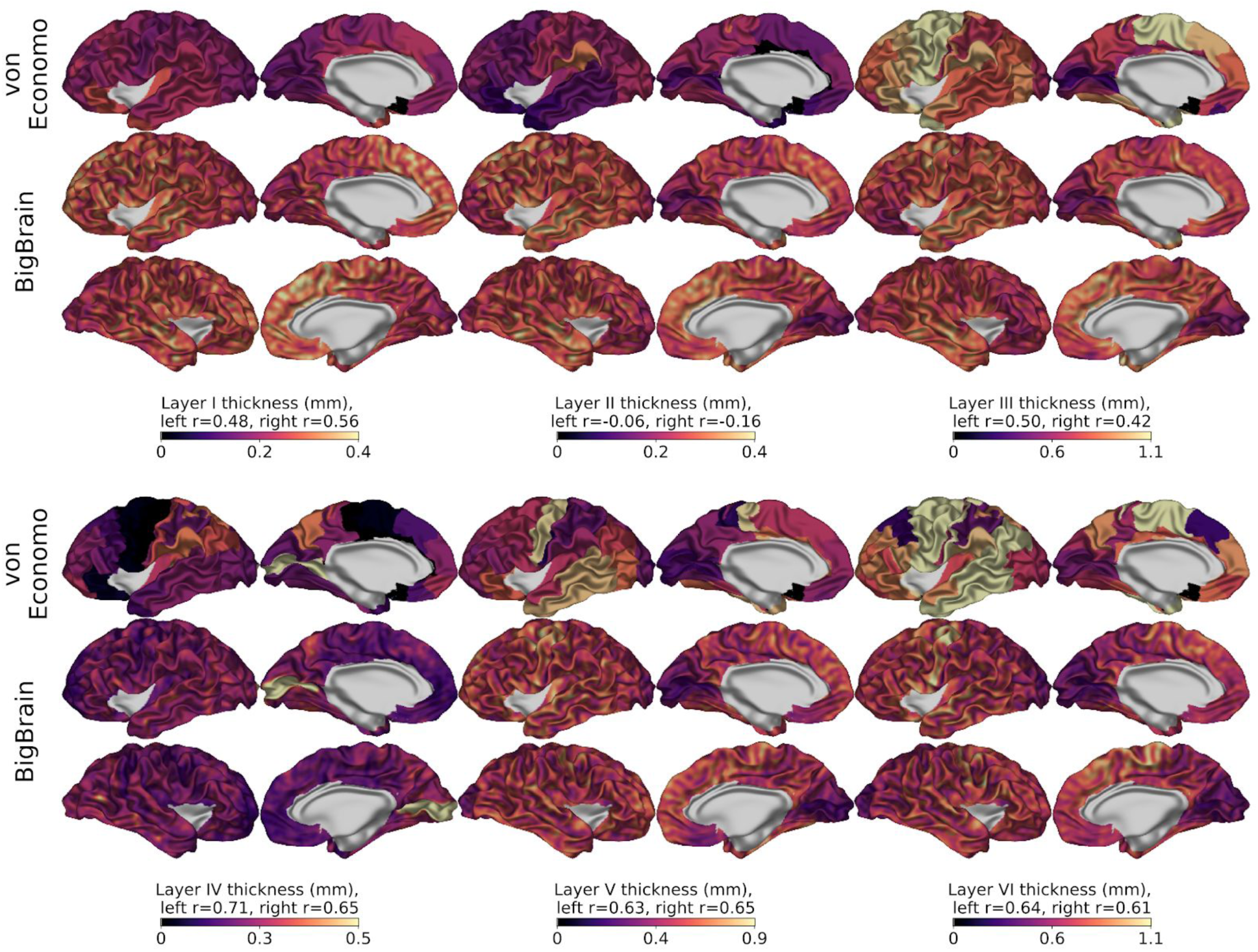
Comparison of von Economo’s laminar thickness maps (coregistered with and visualized on the BigBrain) with laminar thicknesses of the BigBrain for left and right hemispheres. BigBrain thickness values were smoothed across the surface with a 3mm FWHM Gaussian kernel. Layer thickness values strongly correlated between BigBrain and von Economo for all layers except layer II (see Results). Similarities include the clear changes in thickness in pre- and post-central thicknesses of layers III, V and VI. For layer IV, the most striking feature is the abrupt change in layer IV thickness at the V1-V2 border. This abrupt change and the unique features of layer IV in V1 lead us to conclude that the neural network may have internally learned to recognise V1 and apply the appropriate laminar segmentation rules.

Of interest is the clear boundary exhibited by layer IV at the boundary between V1 and V2 in the occipital cortex. This change in thickness is abrupt enough to generate an automated anatomical label for V1 (Figures 2 and 4).

### Cortical gradients and processing hierarchies

Cortical thickness was positively correlated with geodesic distance in visual (left, r=0.56, p=0, right, r=0.42, p=0; von Economo r=0.72, p=0.02), somatosensory (left, r=0.20, p=0, right, r=0.30, p=0; von Economo r=0.64, p=0.12), and auditory cortices (left, r=0.20, p=0, right, r=0.13, p=0; von Economo r=0.92, p=0.01) (Figure 5A-C). By contrast in the motor cortex, thickness was negatively correlated with geodesic distance (left, r=-0.25, p=0, right, r=-0.16, p=0; von Economo r=-0.84, p=0) (Figure 5D). These results are consistent with MRI thickness findings in sensory gradients but contradictory for the fronto-motor gradient.

**Figure 5.**
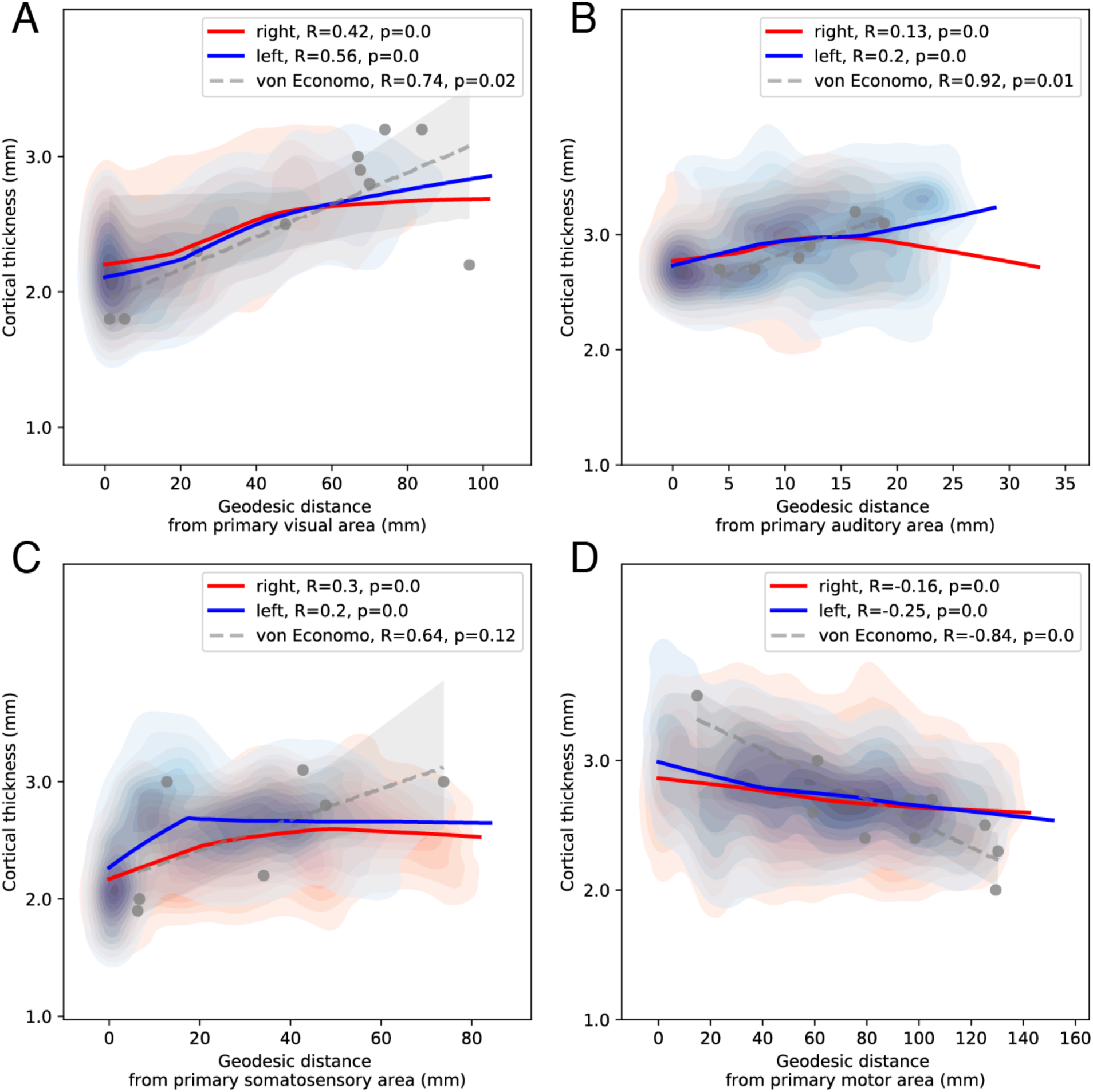
Cortical thickness with increasing geodesic distance from the primary area. To aid visualisation, Locally Weighted Scatterplot Smoothing lines are fit for each hemisphere. For primary visual, auditory and somatosensory cortices (A-C), consistent with MRI studies of cortical thickness, thickness increased with geodesic distance from the primary sensory areas. These trends were also present in the von Economo dataset, where statistical power was limited by the small number of samples. For the motor cortex (D), a negative relationship was present with thickness decreasing from the primary motor cortex into the frontal cortex in the BigBrain dataset and von Economo. This structural gradient is the inverse of the pattern of myelination and of previously reported MRI frontal thickness gradients, but consistent with patterns of structural type and neuronal density. These findings suggest the presence of distinct but overlapping structural hierarchies.

Cortical layers did not contribute equally to the total thickness gradient in the visual and somatosensory cortices (Figure 6A). Layers III and V had the largest contributions to the total thickness gradient, followed by layer VI and then II. A similar but less pronounced pattern was seen within the auditory cortex. In the motor cortex the inverse was true, with decreases in layers III, V and VI. Changes in the same cortical layers appeared to drive gradients in the von Economo laminar thickness measurements (Figure 6B), but due to the small number of recorded samples the confidence intervals were larger and generally included zero.

**Figure 6.**
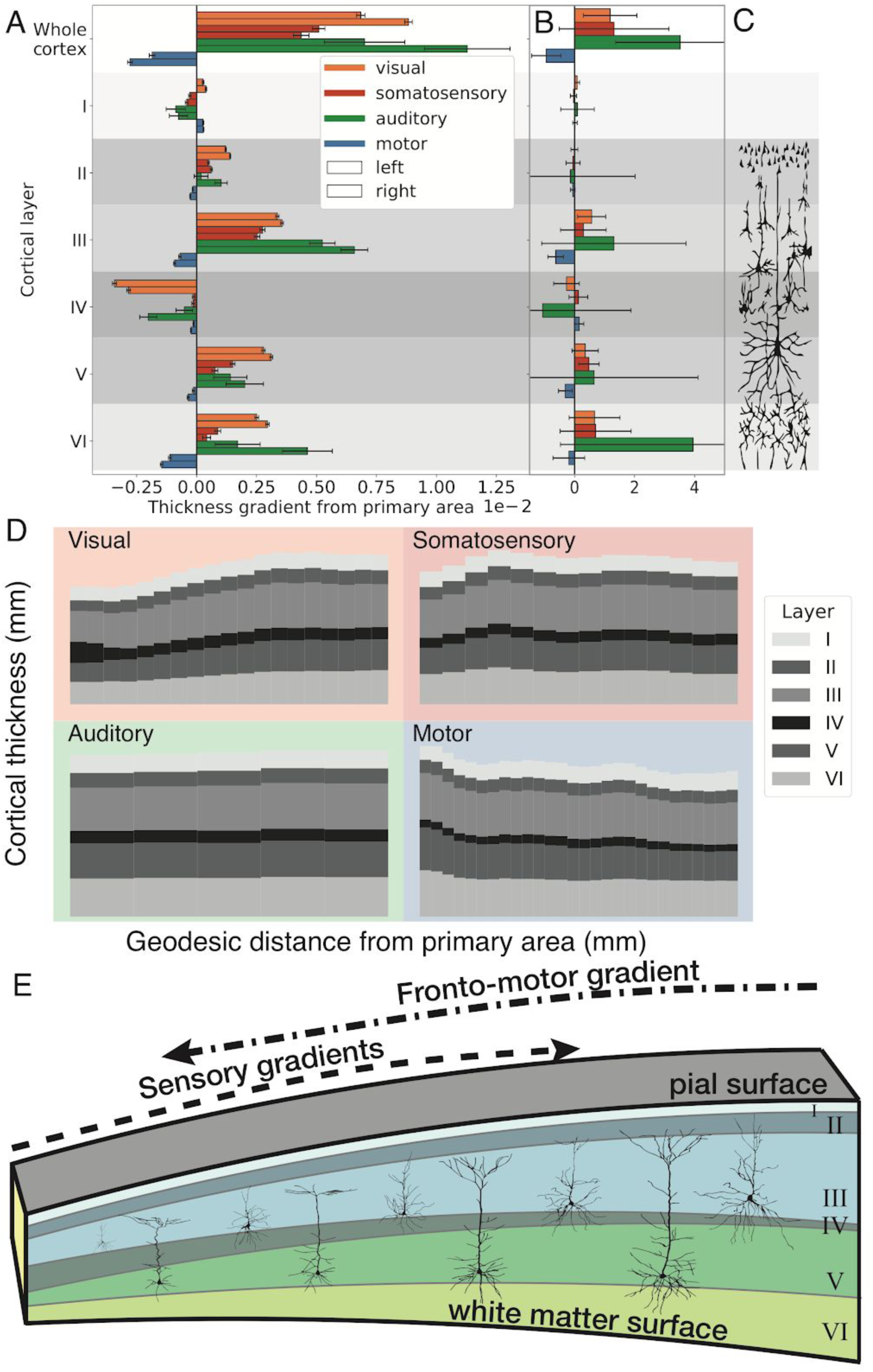
Gradients of cortical and laminar thickness against geodesic distance from primary areas. A) Fronto-motor gradients show an inverse relationship from sensory gradients both on cortical and laminar thicknesses. Increasing sensory cortical thickness gradients were generally driven by thickness increases in layers III, V and VI. By contrast, fronto-motor cortical thickness gradients exhibited decreases in thickness of the same layers. B) The same trends were evident in the von Economo dataset, however due to the small number of recorded samples the confidence intervals were larger and generally included zero. C) Typical neuronal types and morphologies of individual cortical layers. Cortical thickness gradients in either direction are primarily driven by changes in pyramidal cell layers (in layers III, V and VI). D) Layer thicknesses averaged across vertices in a sliding window of geodesic distance values from the primary area for the visual, somatosensory, auditory and motor systems. The motor cortex exhibits the inverse pattern of change to those observed in sensory gradients. E) Single-cell morphological studies of pyramidal neurons in macaque sensory processing pathways reveal increasing dendritic arborisation [65] consistent with the hypothesis that laminar volume changes and ultimately thickness changes represent increases in intracortical connectivity.

Thus, somatosensory and visual areas exhibited positive histological thickness gradients primarily driven by layers III, V and VI. By contrast the fronto-motor areas exhibited an inverse gradient, peaking in the motor cortex and driven by the same layers (Figure 6B & C).

### Neural network training

In the cross-validation, average per-point accuracy on the test fold was 83% ± 2% prior to post-processing, indicating that the network was able to learn generalisable layer-specific features and transfer them to novel cortical areas. The predictions of the model trained on the full dataset were used to create a 3D segmentation of the cortical layers in both hemispheres of the BigBrain dataset (Figure 1).

### Confidence results

Layer confidence maps, given by the difference between prediction values (between 0 and 1) of the highest and second-highest predicted classes for each point, give an approximation of the reliability of laminar segmentations for the cortex where ground truth manual segmentations have not been carried out (Supplementary Figure 2). Throughout the cortex, the network has high confidence for supra-pial and white matter classes. Cortical layers also exhibit consistent confidence maps, with slightly lower confidence for layer IV. This pattern matches with visual observations that layer IV is often the most difficult layer to identify.

### Resolution results

Downsampling BigBrain to decrease the resolution, from 20μm down to 1000μm, progressively decreased the accuracy of the network on the test folds from 85% to 60% (Supplementary Figure 3). However, at 100μm (the approximate upper limit for current high-resolution structural MRI), profiles had sufficient detail to maintain an accuracy of 76%.

### Open data availability

All code used for the segmentation and analysis of cortical layers are being made available at https://github.com/kwagstyl. BigBrain volumes, layer surfaces and intensity profiles are available to download at ftp://bigbrain.loris.ca/ as well as at https://bigbrain.humanbrainproject.org.

## Discussion

We automatically segmented the six histological layers of the cerebral cortex in the 3D reconstructed BigBrain. This is the first whole brain quantitative, laminar atlas with high precision, and the first ever in 3D. Our approach overcomes many historical problems with histological thickness measurements, and provides a higher level of precision and detail than any past laminar atlases. We used this atlas to test for gradients of cortical and laminar thickness within sensory and motor processing hierarchies. Consistent with previous findings using *in vivo* MRI [10] and 2D histological measurements [8], sensory hierarchies exhibited a gradient of increasing cortical thickness from primary sensory to higher order areas. In visual, somatosensory and auditory cortices, these gradients were primarily driven by layers III, V, VI. By contrast, the fronto-motor cortices exhibited a decreasing cortical thickness gradient away from the primary motor cortex towards higher order frontal areas, which was driven by decreases in these layers. These findings highlight the utility of the BigBrain for linking micro- and macro-scale patterns of cortical organisation.

Gradients of thickness are large scale markers of systematic changes in the cortical microcircuit. The volume of the cortex is 80-90% neuropil [33–37], of which 60% is axons and dendrites and the remainder is synaptic boutons, spines and glia. As neuronal density decreases with increasing cortical thickness [38,39], and most of the volume of the cortex is neuropil, increased thickness is most likely to mark increased intracortical connectivity [7]. At a laminar level, the strongest contributors to the overall thickness gradients were layers III, V and VI (Figure 6). Cell-morphological studies in macaques have shown that the cell size and dendritic arborisation of layer III and V pyramidal neurons increase along the visual pathway [18,40,41]. Similarly, afferent axonal patch sizes scale with pyramidal neuronal arborisation [42]. Increasing dendritic arborisation, axonal field size and number of synapses all contribute to an increase in the volume of laminar neuropil and are therefore a likely driver of the laminar and overall thickness gradients measured here. Layer VI also increased in thickness in sensory pathways. However, while neurons located in these layers might exhibit increases of their associated neuropil, the measured thickness change may in part be due to the extended arborisations of III/V pyramidal neurons forming connections within these layers. Histological gradients of layer thickness provide us with a meso-scale link between *in vivo* patterns of MRI cortical thickness and microstructural, neuronal-scale changes in the cortical microcircuit. Such links help us to understand the neurobiological significance of interindividual, longitudinal and neuropathological biomarkers [7].

In contrast to *in vivo* studies of fronto-motor functional, myelin and MRI cortical thickness organisation, which place the primary motor cortex at the same level as primary sensory areas [15,17,27], we found that total and laminar fronto-motor thickness gradients were the inverse of those measured in sensory cortices. This places the fronto-motor cortex in a distinct hierarchical position. Histologically, the motor cortex was especially thick and the thickness decreased with geodesic distance from the primary motor cortex, with layers III, V and VI following a similar inverse pattern. This finding is consistent with reported trends in other histological properties such as laminar structural type [8] and neuronal density [38], as well as the observation that the motor cortex has large, pyramidal neurons with extensive dendritic arborisation [43,44]. It is also in agreement with the distribution of neurotransmitter receptors. The molecular architecture as estimated by neurotransmitter receptors also provides evidence that primary visual and motor cortex are on opposite positions in cortical hierarchy – the acetylcholinerigc M2 receptor, but also NMDA, GABAA, GABAA/BZ, M2, α2, 5-HT2, and D1 receptors show high densities in the primary sensory areas, lower densities in association areas and the primary motor cortex among the lowest [25].

Functionally, these structural differences might be considered in terms of narrow, specific columnar receptive fields for accurate sensory perception [45], and wider receptive fields [46] for the coordination of multiple muscle groups [47] in precise motor control. Such microstructural trends are likely to be a result of matching gradients of genetic expression [48], and may indirectly relate to other microstructural trends including the relative somal size and connectivity patterns of pyramidal neurons in layers III and V [22,49]. Thus there is a coherent group of cortical microcircuit properties which diverges from patterns of cortical myelination and fMRI-derived gradients, establishing the motor cortex at the peak of a gradient of increasing cortical thickness, layer III, V and VI thickness and pyramidal neuronal arborisation, with primary sensory areas at the opposite extreme.

### Atlas of cortical layers

The layers we have generated to test gradient-based hypotheses have applications beyond the scope of this study. Surface-based models of layer structure also create a framework for translating between microstructural modalities and surface-based neuroimaging. For instance, layer segmentations can be used to define regions of interest for further detailed analysis, and for associating cortical *in vivo* and *ex vivo* data to the common BigBrain template. Furthermore, current approaches to measuring laminar structure and function *in vivo* rely on prior models of the cortical layers - for example signal-source simulation in MEG [50] or for sampling laminar BOLD signal in fMRI [51]. The whole-brain histological models for areal layer depth provided here, combined with a thorough understanding of how the layers vary with local cortical morphology [28,52,53] will aid such anatomical models.

### Limitations

It is important to acknowledge that the gradients of laminar thickness measured may be affected by limitations in the BigBrain dataset. The first limitation is that the *post mortem* brain was damaged during extraction and mounting. In some areas this resulted in minor shears. This problem was addressed to some extent through the utilisation of non-linear registration techniques. Nevertheless, some shifts in cortical tissue between consecutive sections are present and will affect the accuracy of layer reconstructions. In other areas, the cortex has been torn. Spatial smoothing and the large total number of sample points make it unlikely that these errors are affecting the results. A second limitation is that there is only one BigBrain. Future work will be necessary to establish the interindividual and age-dependent variability in laminar structure, either using other histological BigBrains or with complementary high-resolution MRI imaging approaches.

## Conclusions

Total cortical thickness and thicknesses for each of the six isocortical layers were measured in the BigBrain to explore the histological drivers of MRI-based thickness gradients. Overall, the pattern of thickness in the BigBrain is consistent with histological atlases of cortical thickness, such as that from von Economo and Koskinas (1925). In the visual and somatosensory cortices, an increasing gradient of histological cortical thickness was identified and found to be primarily driven by layers III, V and VI. In the fronto-motor cortex, the inverse pattern was found. These findings provide a link between patterns of microstructural change and morphology measurable through MRI, and emphasise the importance of testing MRI-based anatomical findings against histological techniques. The laminar atlases provide an invaluable tool for comparison of histological and macroscale patterns of cortical organisation.

## Acknowledgements

Parts of this work have received support from Healthy Brains for Healthy Lives, the Avrith MNI-Cambridge Neuroscience Collaboration award, and the European Union’s Horizon 2020 Framework Programme for Research and Innovation under Grant Agreement No. 785907 (HBP SGA2).

## Methods

### Volumetric data preparation

BigBrain is a 20×20×20μm (henceforth described as 20μm) resolution volumetric reconstruction of a histologically processed *post mortem* human brain (male, aged 65). It is available for download at ftp://bigbrain.loris.ca and is used as a reference brain of the Atlases of the Human Brain Project at https://www.humanbrainproject.eu/en/explore-the-brain/atlases/. In order to run computations on this 1TB dataset, the BigBrain was partitioned into 125 individual blocks, corresponding to five subdivisions in the x, y and z directions, with overlap. The overlap of blocks was calculated to be sufficient such that each single cortical column could be located in a single block, enabling extraction of complete intensity profiles between pairs of vertices at the edge of blocks without intensity values being altered by boundary effects when the data were smoothed. Blocks were smoothed anisotropically [54], predominantly along the direction tangential to the cortical surface, to maximise interlaminar intensity differences while minimising the effects of intralaminar intensity variations caused by artefacts, blood vessels and individual neuronal arrangement [28]. The degree of anisotropic smoothing is determined by repeatedly applying the diffusive smoothing algorithm. The optimal level of smoothing was previously determined and gave an effective maximum full width at half maximum (FWHM) of 0.163mm [28]. For subsequent analyses both the raw 20μm and anisotropically smoothed blocks were used.

Lower resolution volumes were extracted by subsampling the raw BigBrain 20μm volume at 40, 100, 200, 400 and 1000μm. Anisotropically smoothed volumes were also generated at each of these resolutions.

### Profile extraction

Pial and white surfaces originally extracted using a tissue classification of 200μm were taken as starting surfaces [30]. Prior to intensity profile extraction, the start and end surfaces were altered to address several issues. First, the vectors connecting white and grey vertices were altered in order to improve their approximation of columnar trajectories and to minimise intersecting streamlines. Second, “pial” and “white” surfaces were respectively expanded beyond their original limits, extending the extracted profiles to contain the whole cortex with additional padding. This was to enable the network to adjust surface placement of these borders according to features learned from the manual delineation of these boundaries. To achieve this, the following steps were taken:

1. A mid-surface was generated that was closer to the pial surface in sulci and closer to the white surface in gyri, weighting the distance vector by the cortical curvature. Thus the mid-surface was closer to the surface with the higher curvature.
2. This mid-surface was upsampled from 163842 to 655362 vertices to increase its resolution.
3. For each vertex, the vector between nearest points on the pial and white surfaces was calculated.
4. To avoid crossing profiles which can result in mesh self-intersections, the vector components were smoothed across the mid-surface with a FWHM of 3mm (Supplementary Figure 1).
5. Profiles were calculated along these vectors from the mid-surface, extending the profiles 0.5mm further than the minimum distance in the pial or white direction, to ensure the resultant intensity profile captured the full extent of the cortex.

The resultant profiles were less oblique and more likely to be lined up with the cortical columns (Supplementary Figure 1).

Extended intensity profiles were then created by sampling voxels at 200 equidistant points between each pair of vertices from the raw and anisotropically smoothed BigBrain volumes, at each available resolution. For an extended profile of approximately 4mm, this gives a distance of 0.02mm or 20μm between points, corresponding to the highest resolution volume available. To account for the rostrocaudal gradient in staining intensity enabling the network to better generalise between profiles, profile intensity values were adjusted by regressing between mean profile intensity and posterior-anterior coordinate in 3D space.

### Training data

Manual segmentations of the 6 cortical layers were created on 51 regions of the cortex, distributed across 13 of the original histological BigBrain sections at rescanned at a higher in-plan resolution of 5μm (Figure 2). These regions were chosen to give a distribution of examples demonstrating a variety of cytoarchitectures, in both gyri and sulci, from sections where the cortex was sectioned tangentially. Layers were segmented according to the following criteria. Layer I, the molecular layer, is relatively cell-sparse with few neurons and glia. Layer II, the external granular layer, is a much more dense band of small granular cells. Layer III, the external pyramidal layer, is characterised by large pyramidal neurons that become more densely packed towards its lower extent. Layer IV, the internal granular layer (usually referred to simply as the “granular layer”), generally contains only granular neurons, bounded at its lower extent by pyramidal neurons of layer V. Layer V, the internal pyramidal layer, contains large but relatively sparse pyramidal neurons while layer VI, the multiform layer, has a lower density of pyramidal neurons [1]. Segmentations were verified by expert anatomists: SB, NPG and KZ. This resolution is sufficient to distinguish individual cell bodies, a prerequisite to analyze their distribution pattern in cortical layers and to delineate the layers. Averaged across all training examples, layer classes contributed to profiles as follows: background/CSF (cerebrospinal fluid): 14.6%, layer I: 7.5%, layer II: 5.6%, layer III: 20.8%, layer IV: 5.5%, layer V: 14.8%, layer VI: 17.8%, white matter: 13.4%. For the cortical layers, these values represent an approximate relative thickness.

Manual segmentations were then co-registered to the full aligned 3D BigBrain space. The manually drawn layers were used to create corresponding pial and white surfaces. These cortical boundaries were extended beyond layer VI and beyond the pial surface between 0.25mm and 0.75mm so as to match the variability of cortical extent in the test profile dataset. Training profiles were created by sampling raw, smoothed and manually segmented data, generating thousands of profiles per sample. Each pixel in the labelled data had a class value of 0-7, where pixels superficial to the pial surface were set to 0, followed by layers numbered 1-6, and white matter was classed as 7. This 1D profile-based approach greatly expanded the training dataset from 51 labelled 2D samples to 564,041 profiles.

### Neural network

A 1D convolutional network for image segmentation was created to enable the identification of laminar-specific profile features which can appear at a range of cortical profile depths [28]. The network was created using stacked identical blocks. Each block contained a batch normalisation layer to normalise feature distributions between training batches, a rectify non-linearity layer used to model neurons which can have a graded positive activation but no negative response [55], and a convolutional layer [56]. There was a final convolutional layer with filter size 1 and 8 feature maps, one for each class. The cost function was median class-frequency weighted cross-entropy. Class-frequency weighting was added to weigh errors according to the thickness of the layers so that incorrectly classified points in thinner layers were more heavily weighted than errors in incorrectly classified thicker layers [57]. Raw and smoothed profiles were considered as two input channels. The network was iteratively trained until the accuracy did not improve for 50 epochs (all training profiles are seen once per epoch). At this point, the previous best model was saved and used for subsequent testing on the full dataset. When testing the network, a soft maximum was then applied to detect the most likely layer class for each point. The output was a matrix of 8 (predicted layers) by 200 (sample points) by 655362 (vertices on a mesh) by 2 (cortical hemispheres).

For each vertex, a measure of confidence was calculated from these predictions. Per point confidence is the difference between the prediction value for the highest predicted class and the value of the 2nd highest predicted class. Per class/layer confidence is the mean confidence for all points in that class/layer. The per-vertex summary measure is the mean across all points in the profile. These measures give an indication of the relative confidence for the regional and laminar classifications.

### Hyperparameter optimisation and cross-validation

Here, a set of 50 experiments with random hyperparameters was carried out to explore their impact on training accuracy (there is no consensus method for finding optimum parameters for a neural network). Learning rate, convolutional filter size, number of layers (blocks), weight decay and batch size were all varied. In summary, the final network was initialised with 6 layers, filter size = 49, learning rate = 0.0005, weight decay = 0.001, where the learning rate determines the amount weights are updated on each iteration, and weight decay determines the rate at which weights decrease each iteration (which helps prevent overfitting).

For network cross validation, the manually labelled areas were subdivided into 10 equally sized random subsets or folds. Initially, 2 folds were removed from the dataset during training and network weights were optimised for segmenting samples on one of these folds. This trained network was then used to predict layers on the final previously unseen test fold from which the accuracy was calculated. This process was repeated 10 times to generate an estimate of the network’s ability to segment novel cortical regions. The same process was carried out using profiles extracted at all available resolutions.

For generating BigBrain layer segmentations, the network was trained on the full training dataset and tested on all intensity profiles.

### Shrinkage estimate

Histological processing, including fixation and sectioning, causes distortion of the tissue that is non-uniform in the x,y and z directions. Part of this distortion was corrected in the original reconstruction of the BigBrain [16].

Initial shrinkage of the brain during fixation prior to sectioning was calculated based on the estimated fresh volume of BigBrain, inferred from the original fresh weight, and the volume after histological processing. This gave a volume-based (3D) shrinkage factor of 1.931, which corresponds to an isotropic length-based (1D) shrinkage factor of 1.245.

To estimate the scale of shrinkage in each of the three orthogonal directions, the BigBrain volume was linearly coregistered to a volumetric MRI template derived from a group of older subjects (ADNI) [58]. The transformation matrix gave linear scale factors of 1.15, 1.22 and 1.43 in the x,y and z directions, with a mean of 1.26. The concordance of these measures of shrinkage suggests that subsequent thickness and length estimates can be adequately corrected for comparison to *in vivo* measures.

Thus, to approximately compensate for the non-uniform compression of xyz, we transformed the mesh surfaces into MNI space based on the ADNI template. Subsequent thickness analyses were carried out on the transformed meshes. Such compensation for shrinkage is necessary when analysing cortical thickness gradients on oblique profiles in 3D over the whole brain. Non-linear corregistration was not applied as this can lead to localised warping and non-biological thickness measurements.

### Surface reconstruction: post-processing 1D profiles

1D classified profiles were transformed into mesh layer boundary reconstructions as follows. Transitions between predicted layers were located for each profile and the coordinates of these transitions became vertex locations for the new layer meshes. For the small number of vertices where the network failed (less than 1%), vertex locations were interpolated from the neighbouring vertices. Surface indices were smoothed 0.5mm FWHM across the cortical surface and 20 iterations of shrinkage-free mesh smoothing was applied to the output surface [59]. This removed non-biologically high frequency changes in surface curvature, most commonly due to minor, local misalignment of consecutive 2D coronal sections.

### Cortical thickness, layer thickness

Cortical thickness was calculated between pial and white cortical surfaces and laminar thicknesses were calculated between adjacent pairs of cortical surfaces.

### Masking

Manual masks were created to remove the medial wall, the allocortex, including parts of the cingulate and entorhinal cortex which do not have 6 layers, and large cuts in the anterior temporal cortex (caused by the saw during extraction of the brain from the skull) from subsequent analyses.

### Surface-based parcellations

For comparison, several existing surface-based parcellations defined upon *in vivo* human template surfaces (von Economo: [8,60]; Glasser: [61]) were coregistered to the BigBrain cortical surfaces using an adaptation of the Multimodal Surface Matching approach [62,63].

### Gradients and processing hierarchies

Surface labels for the primary visual, auditory, somatosensory and motor areas were manually delineated on each hemisphere using morphological markers and histological characteristics (Supplementary Figure 5a). For each system, a larger area containing associated cortical regions was manually delineated (Supplementary Figure 5b) [10,17,61]. For each vertex within the associated cortical regions, geodesic distance from the primary sensory labels was calculated (Supplementary Figure 5b) [10,17,61,64].

## Supplementary Information for

**Supplementary Figure 1.**
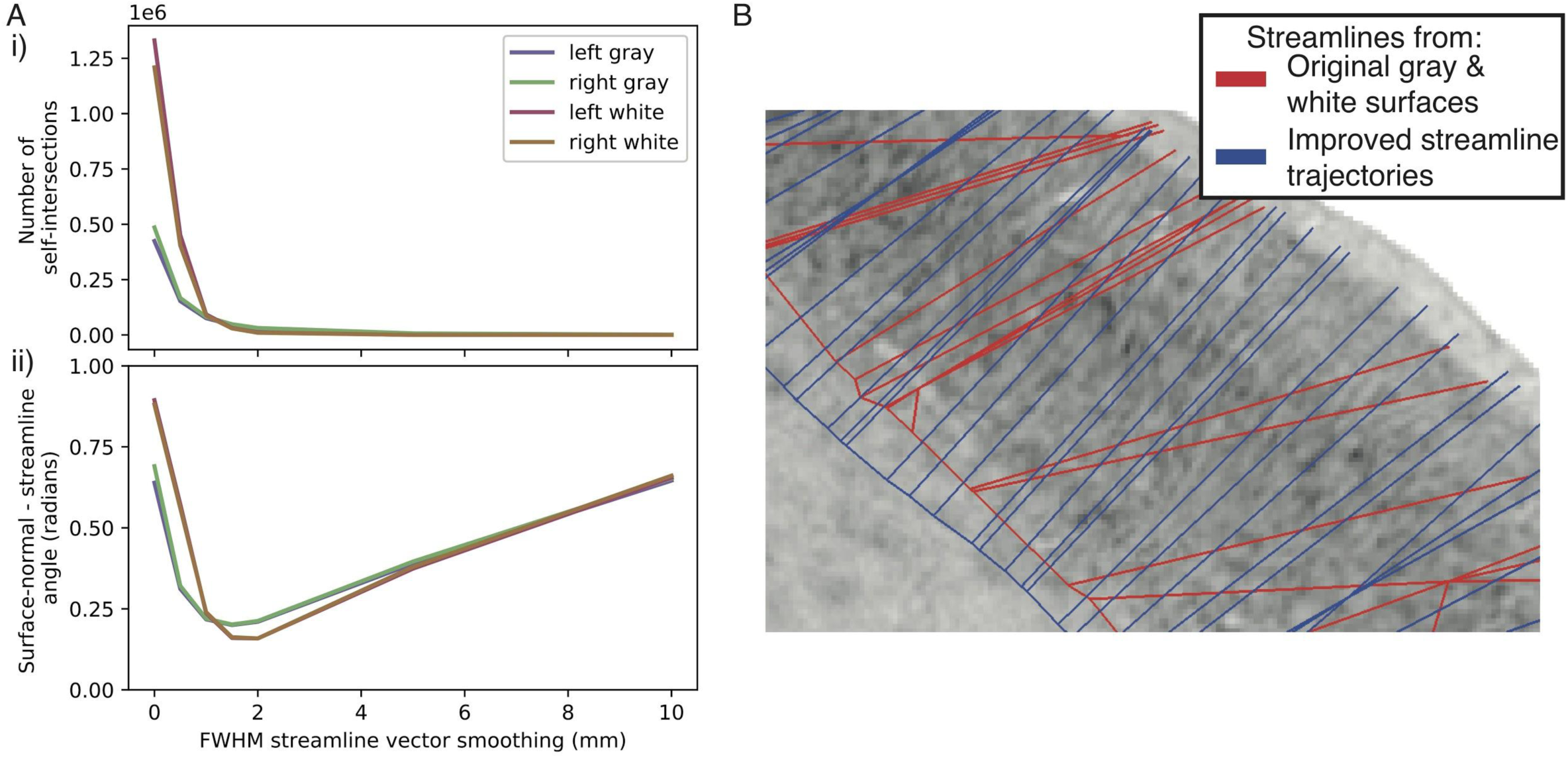
Improving streamline trajectories. A) Streamline vectors were smoothed across the cortical surface by varying degrees to assess the impact of smoothing on i) the number of self-intersections in the pial and white surfaces and ii) the angle between the streamline and the normal vector on the pial and white surfaces. These optimisation curves demonstrate that a FWHM of around 2mm drastically decreases the number of self-intersections and obliqueness of the streamline vectors relative to the pial and white surfaces. B) Visualising streamlines against a histological section. Streamlines more closely follow visible cortical columnar trajectories after this improvement (blue) relative to before this streamline vector smoothing process (red).

**Supplementary Figure 2.**
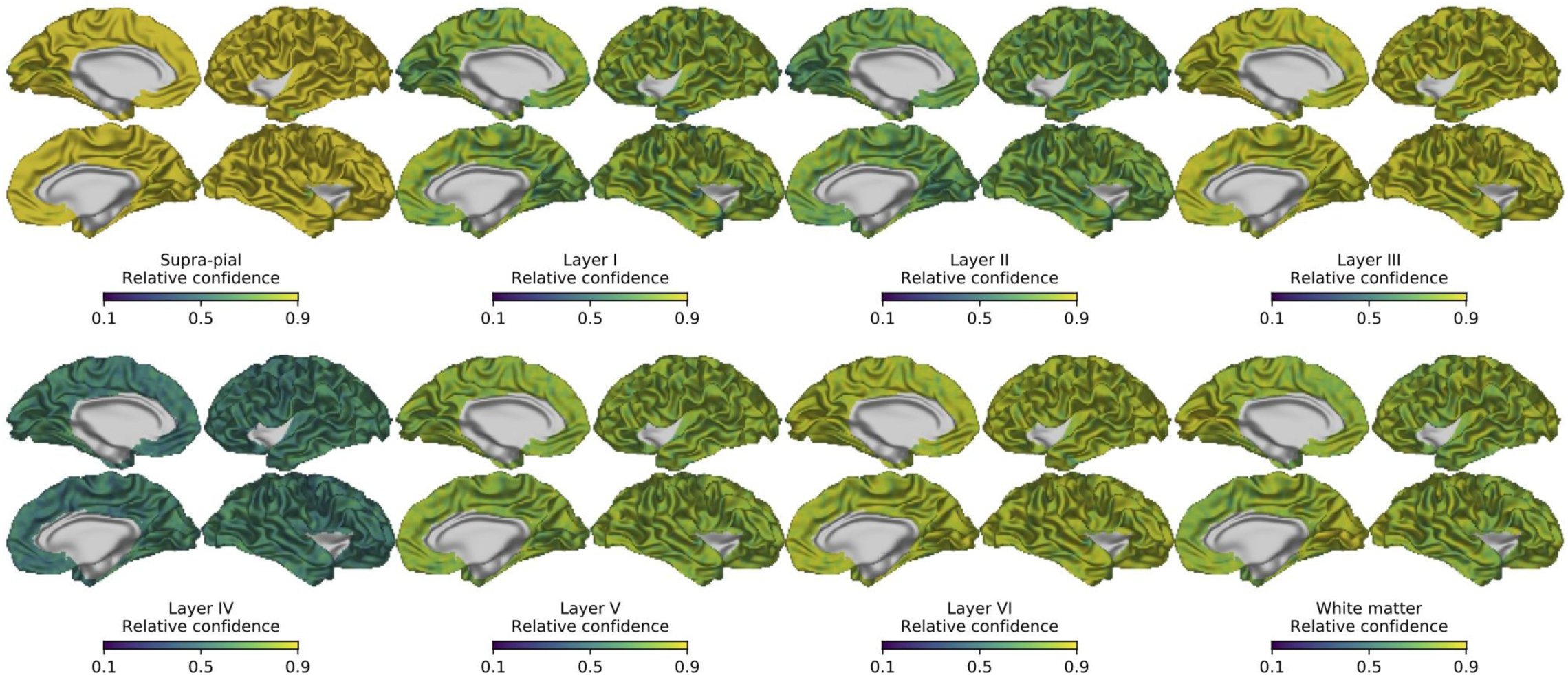
Layer confidence maps. Per-vertex confidence is defined as the difference between the prediction value for the highest predicted class and the value of the 2nd highest predicted class, averaged over the whole profile. This gives an approximation of the reliability of laminar segmentations for the cortex where ground truth manual segmentations have not been carried out. Confidence for supra-pial and white matter classes was high throughout the cortex, thus increasing the confidence in overall cortical thickness measures. Layers exhibit relatively consistent confidence maps, with layer IV least confident overall. This pattern matches with visual observations that layer IV is the most difficult to identify. Regional variations in confidence can guide the choice of target regions for future extensions to the training data.

**Supplementary Figure 3.**
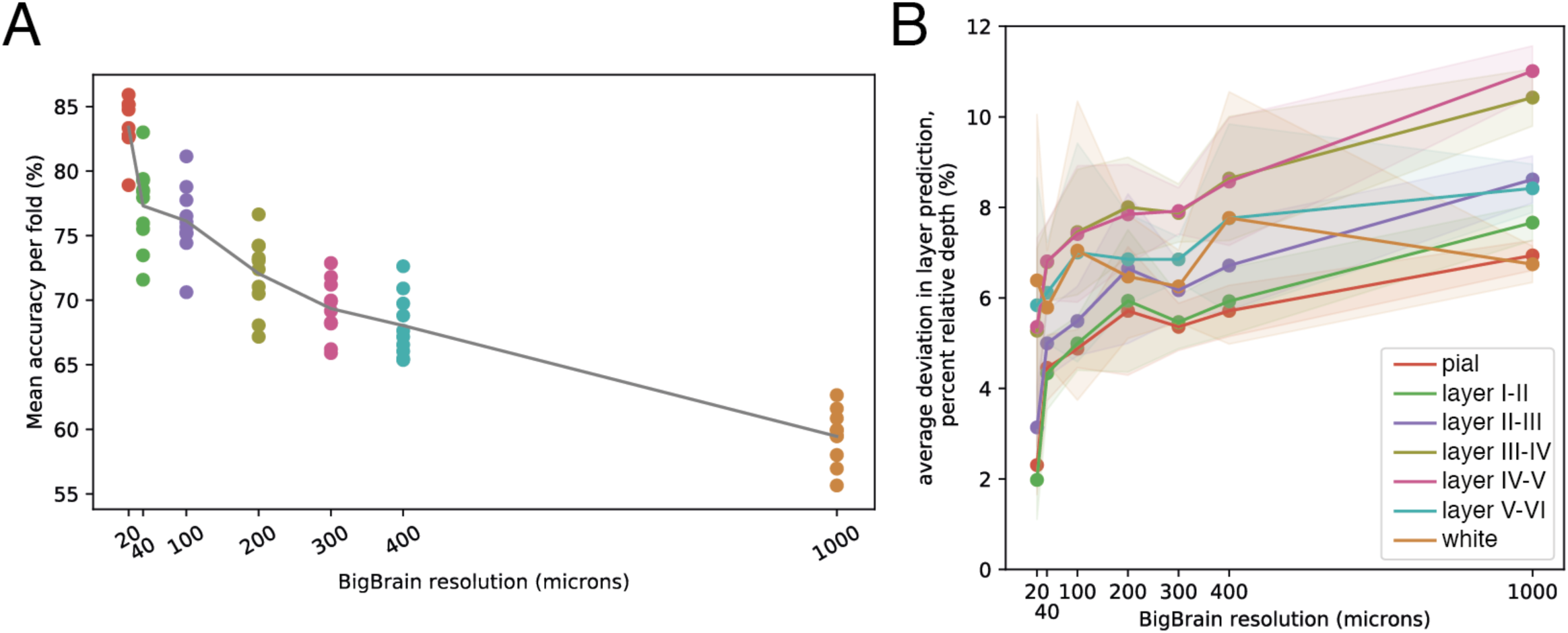
Impact of voxel resolution on overall and layer accuracies. A) Overall per-point accuracies on withheld test regions calculated using 10-fold validation. Accuracy decreases with decreasing resolution. B) Mean deviation in depth prediction on test folds between prediction and manually defined layers. Pial/layer I and layer I-II boundaries exhibited the smallest deviations, followed by II/III, with layer III/IV and VI/white boundaries exhibiting larger deviations.

**Supplementary Figure 4.**
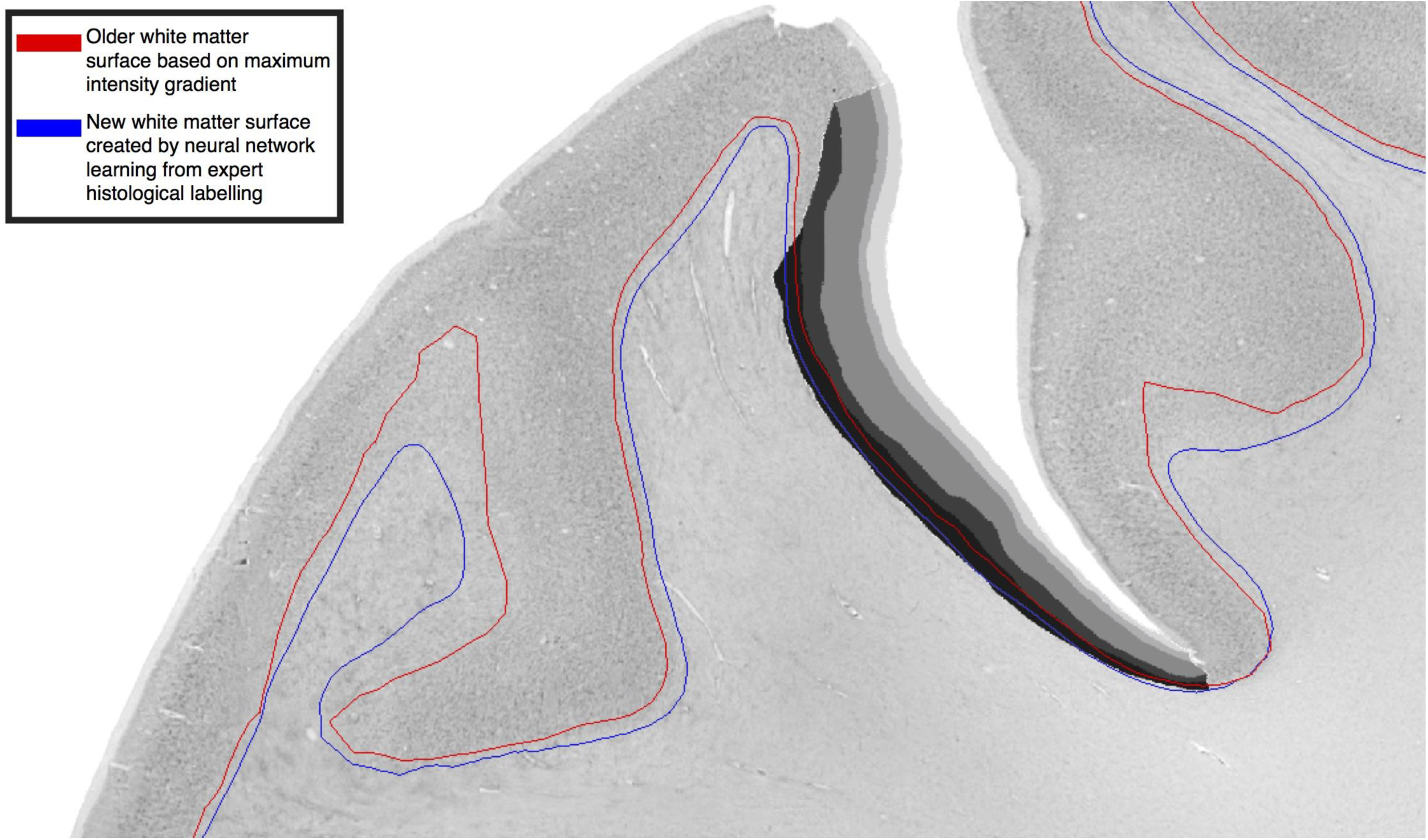
Comparison of white matter surfaces generated by the neural network and by placing the white surface at the maximum intensity gradient. For visual comparison, the surfaces are overlaid on a 2D section, where manually segmented layers are available. The maximum intensity gradient white surface (red) was identified on lower-resolution data (200μm). This surface was consistently superficial to the new surface (blue), which was created based on features derived from the histological definition of the white matter surface, which is determined by the absence or presence of cortical neurons. The blue surface more consistently follows the layer VIa/VIb boundary. This systematic difference highlights the role of using histological expertise when translating across scales and fields to ensure consistent definitions. It also raises an important question on the placement of the white surface in MRI cortical reconstructions, which is placed at the maximum MRI intensity gradient. This gradient is determined predominantly by myelin contrast, and therefore influenced by changes interregional and longitudinal in cortical myelination. Future cortical segmentation algorithms need to be developed with closer reference to histological definitions of the gray/white boundary.

**Supplementary Figure 5.**
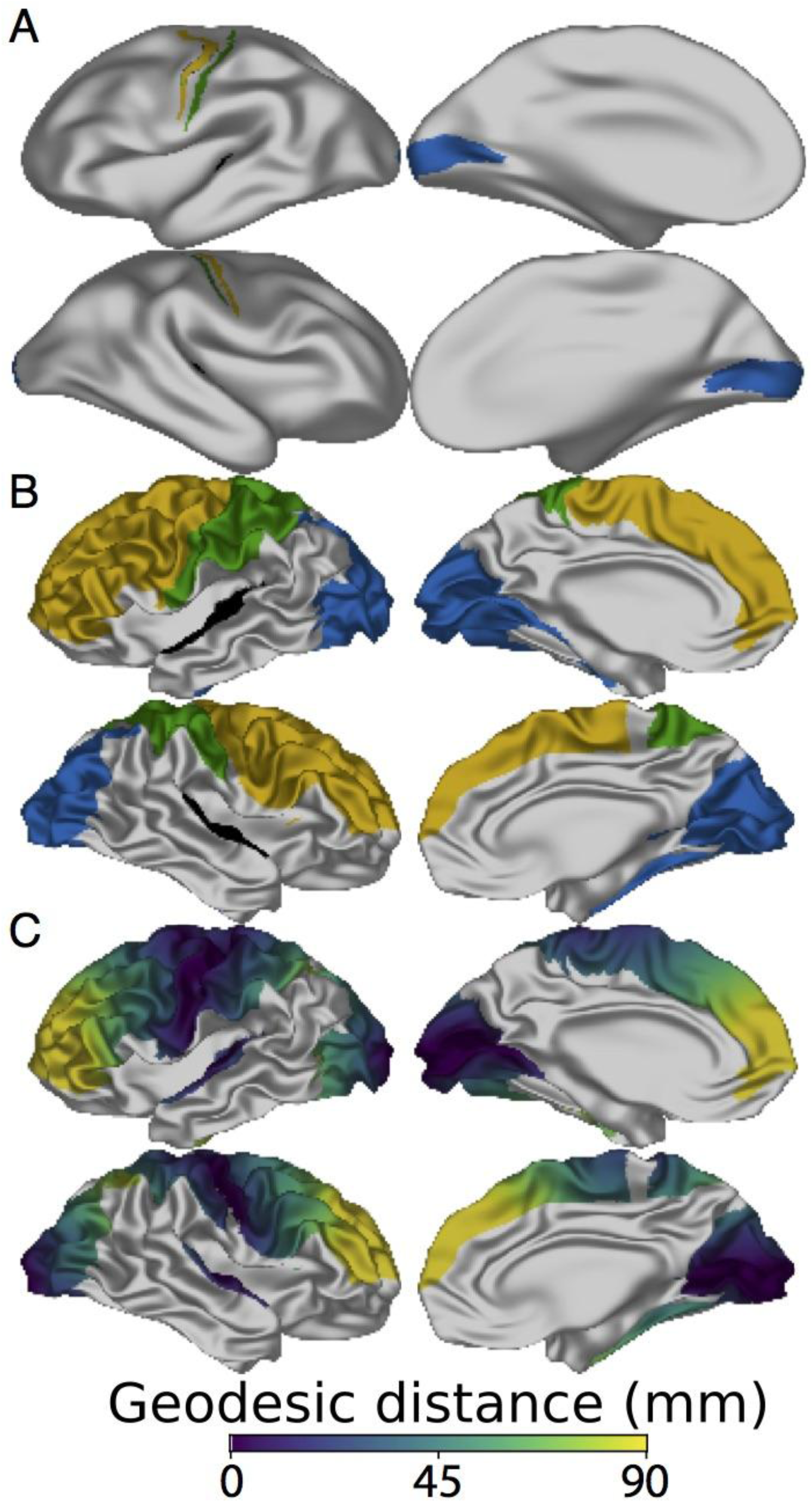
A. Manually segmented primary visual (blue), primary auditory (black, partially buried in the lateral sulcus), primary somatosensory (green) and primary motor (yellow) areas, projected onto a heavily smoothed surface. B. Manually segmented regions across which cortical and laminar hierarchical thickness gradients were calculated. C. Geodesic distance across the cortical surface from the primary areas.

